# Dynamic patterns of cortical expansion during folding of the preterm human brain

**DOI:** 10.1101/185389

**Authors:** Kara E. Garcia, Emma C. Robinson, Dimitrios Alexopoulos, Donna L. Dierker, Matthew F. Glasser, Timothy S. Coalson, Cynthia M. Ortinau, Daniel Rueckert, Larry A. Taber, David C. Van Essen, Cynthia E. Rogers, Christopher D. Smyser, Philip V. Bayly

**Affiliations:** Department of Biomedical Engineering, Washington University in St. Louis, St. Louis, MO, 63130; Department of Computer Science, Imperial College, London, SW7 2AZ, UK; Department of Biomedical Engineering and Department of Perinatal Imaging and Health, Division of Imaging Sciences, St Thomas’ Hospital, Kings College London, SE1 7EH, UK; Department of Neurology, Washington University School of Medicine, St. Louis, MO, 63110; Mallinckrodt Institute of Radiology, Washington University School of Medicine, St. Louis, MO, 63110; Department of Neuroscience, Washington University School of Medicine, St. Louis, MO, 63110; St. Luke’s Hospital, St. Louis, MO, 63017; Department of Pediatrics, Washington University School of Medicine, St. Louis, MO, 63110; Department of Mechanical Engineering and Material Science, Washington University in St. Louis, St. Louis, MO, 63130; Department of Psychiatry, Washington University School of Medicine, St. Louis, MO, 63110

**Keywords:** Cortex, growth, strain energy, registration, development

## Abstract

During the third trimester of human brain development, the cerebral cortex undergoes dramatic surface expansion and folding. Physical models suggest that relatively rapid growth of the cortical gray matter helps drive this folding, and structural data suggests that growth may vary in both space (by region on the cortical surface) and time. In this study, we propose a new method to estimate local growth from sequential cortical reconstructions. Using anatomically-constrained Multimodal Surface Matching (aMSM), we obtain accurate, physically-guided point correspondence between younger and older cortical reconstructions of the same individual. From each pair of surfaces, we calculate continuous, smooth maps of cortical expansion with unprecedented precision. By considering 30 preterm infants scanned 2-4 times during the period of rapid cortical expansion (28 to 38 weeks postmenstrual age), we observe significant regional differences in growth across the cortical surface that are consistent with patterns of active folding. Furthermore, these growth patterns shift over the course of development, with non-injured subjects following a highly consistent trajectory. This information provides a detailed picture of dynamic changes in cortical growth, connecting what is known about patterns of development at the microscopic (cellular) and macroscopic (folding) scales. Since our method provides specific growth maps for individual brains, we are also able to detect alterations due to injury. This fully-automated surface analysis, based on tools freely available to the brain mapping community, may also serve as a useful approach for future studies of abnormal growth due to genetic disorders, injury, or other environmental variables.

**Significance Statement:** The human brain exhibits complex folding patterns that emerge during the third trimester of fetal development. Minor folds are quasi-randomly shaped and distributed. Major folds, in contrast, are more conserved and form important landmarks. Disruption of cortical folding is associated with devastating disorders of cognition and emotion. Despite decades of study, the processes that produce normal and abnormal folding remain unresolved, although the relatively rapid tangential expansion of the cortex has emerged as a driving factor. Accurate and precise measurement of cortical growth patterns during the period of folding has remained elusive. Here, we illuminate the spatiotemporal dynamics of cortical expansion by analyzing MRI-derived surfaces of preterm infant brains, using a novel strain energy minimization approach.

---

**D**uring the final weeks of fetal or preterm brain development, the human cerebral cortex dramatically increases in surface area and undergoes complex folding (Fig. 1). This period represents an important phase of neurodevelopment, as crucial brain regions undergo changes in connectivity and cellular maturation (1, 2). Physical simulations suggest that folding may result from mechanical instability, as the outer gray matter grows faster than underlying white matter (3, 4). Such models accurately predict stress patterns within folds and explain abnormal folding conditions such as polymicrogyria and pachygyria. However, recent models, which consider uniform cortical growth on a realistic brain geometry, do not fully reproduce highly conserved (primary) folding patterns observed in the human brain (4). This suggests a role for other hypothesized factors such as axon tension in white matter (5), regional differences in material properties, or regional differences in growth (6).

**Fig. 1.**
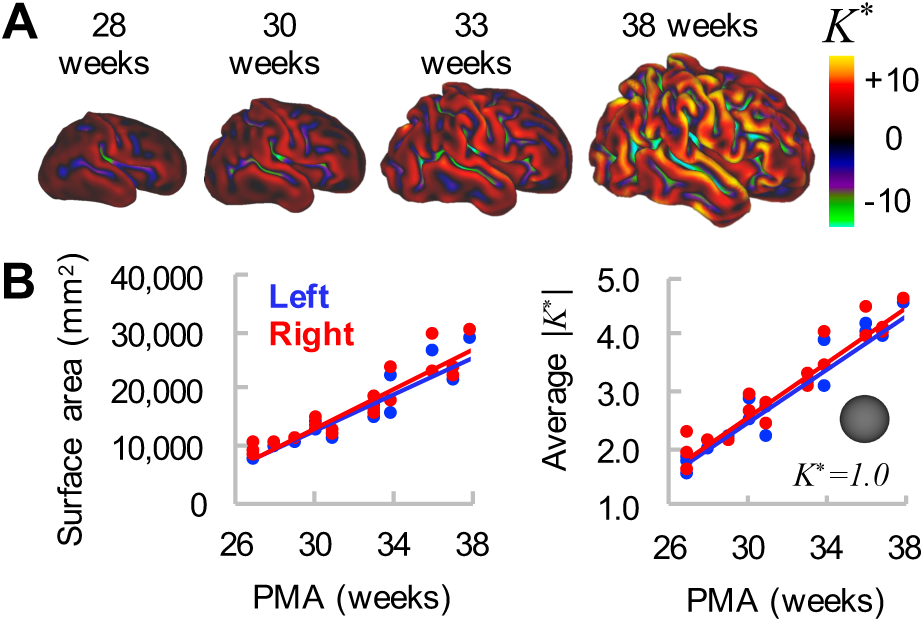
Global measures of cortical surface area and folding increase over time. *(A)* Cortical mid-thickness for individual hemisphere at 28, 30, 33, and 38 weeks PMA. Color overlay represents non-dimensional curvature, *K\**, a useful metric of the degree of folding. *(B)* Total cortical surface area (left) and average magnitude of *K\** (right) increase with time. However, these global measures do not provide information about regional variations in maturation and morphology.

Advances in magnetic resonance imaging (MRI) and cortical reconstruction have enabled detailed quantification of brain structure and connectivity during brain development (7-9). Nonetheless, measuring patterns of physical growth over time presents a unique challenge, as quantification of local growth requires precise identification of corresponding points between multiple scans. To date, clinical studies often rely on global measures of shape or total surface expansion (8) despite evidence of important regional differences (10). Primary sensory and motor regions exhibit earlier maturation of synaptic density and folding than other areas (2, 7). Furthermore, subtle or localized folding abnormalities have been linked to disorders such as epilepsy, autism, and schizophrenia (11-13). To address this issue and attain more detailed measures, several groups have analyzed user-delineated regions of interest (ROIs) (14-16). However, manual definition of ROIs may introduce bias or error and typically requires labor-intensive editing. Furthermore, *a priori* parcellation can lead to skewed or weakened conclusions if the true effect does not fall neatly within the assumed area.

In this study, we employ an ROI-independent approach to estimate spatiotemporal patterns of cortical growth over the course of human brain folding. Harnessing recent developments in *anatomical* Multimodal Surface Matching (aMSM) (17, 18), we achieve physically-guided point correspondence between cortical surfaces of the same individual across multiple time points. This was accomplished by automatically matching common features (gyri and sulci) within a spherical framework and penalizing physically unlikely (energy-expensive) deformations on the anatomical surface, an approach first described in ferret models (19, 20). While alternative methods have been proposed to match anatomical surfaces (21-23), none provide MSM’s flexibility in terms of data matching (17) or utilize penalties inspired by physical behavior of brain tissue (19). Our approach produces smooth, regionally-varying maps of surface expansion for each subject analyzed. By considering right and left hemispheres from 30 preterm subjects, scanned at different intervals from approximately 28 to 38 weeks postmenstrual age (PMA), we observe consistent, meaningful patterns of differential growth that change over time.

This study provides a spatially comprehensive, quantitative analysis of cortical expansion dynamics during human brain folding. We report statistically significant regional differences consistent with established patterns of cellular maturation and the emergence of new folds. These findings, which suggest that prenatal cortical growth is not uniform, may guide future studies of regional maturation and more accurate simulations of cortical folding. Furthermore, since our tools are freely available to the neuroscience community (9, 17, 24, 25), the approach presented here can be widely applied to future studies of development and disease progression.

## Results

To visualize changes in cortical growth over the course of brain folding, we analyzed right and left hemispheres from 30 very preterm infants (born <30 weeks PMA, 15 male, 15 female) scanned 2-4 times before or at term-equivalent (36-40 weeks PMA). Six subjects with injury were excluded from group analysis (see Materials and Methods for criteria), but all were analyzed longitudinally to assess individual growth patterns.

### aMSM produces accurate point correspondence and smooth growth estimation across the cortex for individual subjects

Multimodal Surface Matching uses a flexible spherical framework to align surfaces based on a range of available surface data (17). As shown in Fig. 2, 3D anatomical surfaces (Fig. 2*A*) and corresponding data (eg. univariate patterns of curvature, Fig. 1*A*) are projected to a standard spherical surface in order to provide a simpler geometry for registration (Fig. 2*B*). Spherical registration shifts points on the input sphere until data is optimally aligned with that of the reference sphere (Fig. 2(7), such that reprojection onto the input anatomical surface reveals accurate point correspondence with the reference surface (Fig. 2*D*). In this study, we use mean surface curvature *(K)*, calculated at the cortical mid-thickness, to drive initial matching. To reduce unrealistic distortions induced by both curvature matching and spherical projection, we further refine our registration to minimize physical strain energy (Eq. 1) between the input and reference anatomical surfaces. Unlike other spherical registration methods, which reduce distortions on the sphere, this allows us to explicitly minimize deformations that are energetically unfavorable (and thus unlikely), greatly reducing artifacts associated with the spherical projection process. (See *SI text* for conceptual examples and validation.)

**Fig. 2.**
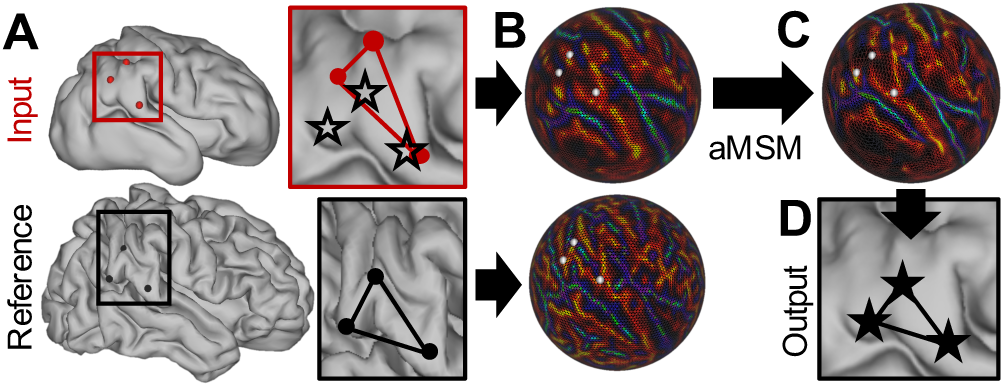
Longitudinal (intra-subject) registration with aMSM. *(A)* Cortical mid-thickness surfaces (ordered sets of vertices in 3D) were generated at multiple time points for each individual. After resampling to a standard mesh, vertices on the input surface (red dots) do not correspond to the same locations on the reference surface (black dots). Red and black boxes outline a region of interest, in which black stars to represent plausible locations for point correspondence. (*B*) Mean curvatures (generated from original topologies, Fig. 1*A*), along with deformations (generated from area-normalized input and reference anatomical surfaces), are projected to a spherical framework to drive registration. (*C*) aMSM moves points on the input sphere in order to (1) optimize curvature matching and (2) minimize deformations between the anatomical surfaces. (*D*) Projection of shifted vertices reveals new anatomical locations with plausible alignment and reduced deformations.

Using aMSM, we were able to obtain physically justified point correspondence and smooth cortical expansion patterns at the individual level. Figure 3 shows results for a representative subject at multiple developmental periods: 27 to 31 weeks, 31 to 33 weeks, 33 to 37 weeks, and directly from 31 to 37 weeks PMA. Qualitatively, plotting the same color map on registered younger (top) and older (bottom) geometries facilitates visualization of the accurate point correspondence for each time period. Quantitative analysis found significant improvements in curvature correlation and strain energy after aMSM registration (Table S1).

**Fig. 3.**
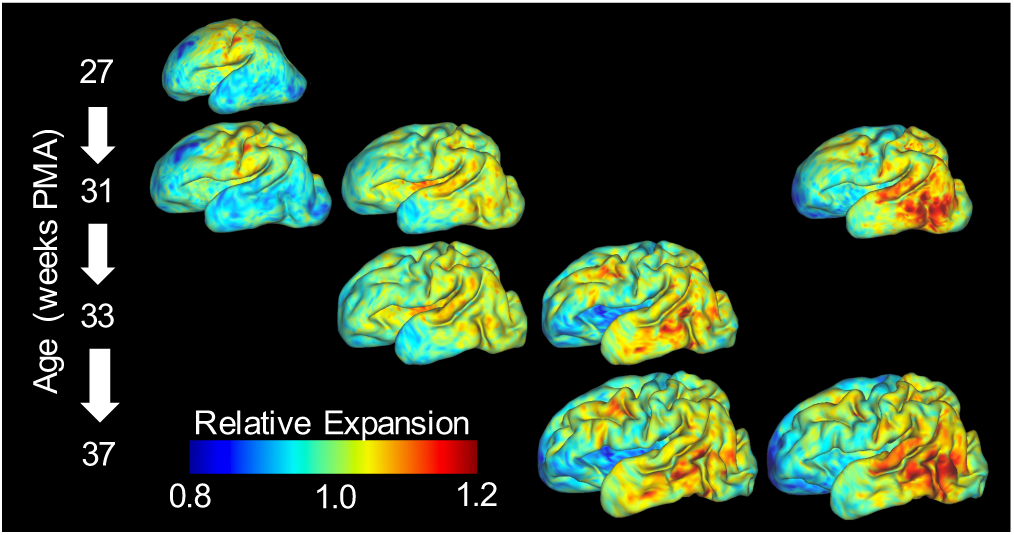
Gradients of cortical expansion are evident in individual brains. Relative cortical expansion (local surface expansion normalized by total hemisphere expansion) is shown for each developmental period. For each column, the same map is overlaid on younger (top) and older (bottom) surface to visualize point correspondence after longitudinal registration. From left to right, relative expansion is estimated for growth from 27 to 31 weeks PMA, 31 to 33 weeks PMA, 33 to 37 weeks PMA, and directly from 31 to 37 weeks PMA. True cortical expansion is equal to relative expansion (plotted) multiplied by global cortical expansion of 1.26, 1.37, 1.70, and 2.33, respectively.

### Spatial patterns of preterm growth are consistent across subjects and correspond to actively folding regions

For comparisons across the entire cohort, individuals were also registered to a 30-week atlas generated from our cohort (Fig. 4*A*, see *SI text* for details). Since folding patterns (primary sulci) are similar across individuals at 30 weeks PMA (7), and because most subjects were scanned near 30 weeks PMA, this atlas served as an appropriate reference for group analyses. Once registration was established between 30-week individual surfaces and the 30-week atlas (inter-subject registration), individual growth metrics (determined by intra-subject registration) were transformed to the atlas, facilitating statistical comparisons across all individuals and time points.

**Fig. 4.**
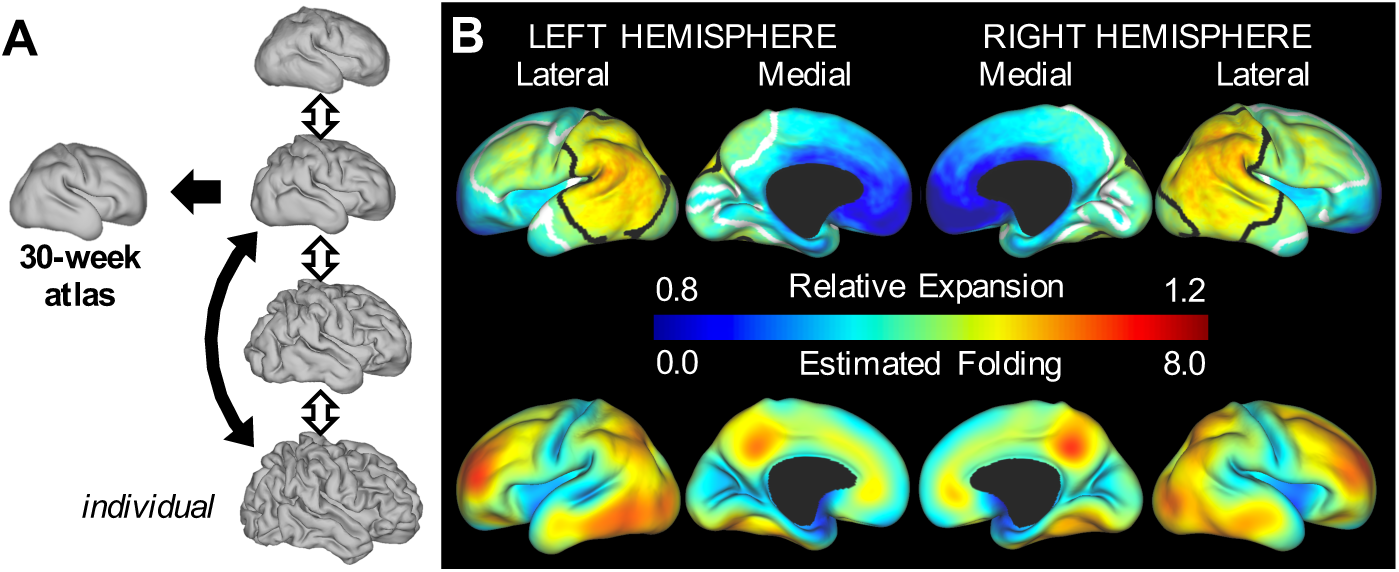
Gradients of cortical expansion and folding are consistent across subjects. *(A)* Correspondence between individual surfaces (closest to 30-weeks, black arrow) and the 30-week atlas was determined by aMSM registration. Using intra-subject correspondence from aMSM (double arrows), individual growth patterns from each period are registered to the 30-week atlas for inter-subject statistics. (S) For the period from 30 to 38 weeks PMA (black double arrow in *A)*, relative area expansion (top) and regional folding (bottom) were highest in the lateral parietal-temporal-occipital region and lateral frontal lobe (n=20). Black and white contours enclose regions where relative expansion is significantly higher and lower than the global average.

To determine whether regional differences in cortical expansion are conserved across individuals, we first considered the period from 30±1 to 38±2 weeks PMA (n=20 subjects without significant injury). Permutation Analysis of Linear Models (PALM) with threshold-free cluster enhancement (25) revealed significantly higher cortical expansion in the lateral parietal, occipital, and temporal regions and significantly lower cortical expansion in medial and insular regions (Fig. 4*B*, top). These patterns were consistent across right and left hemispheres.

At the individual level, we observed highest cortical expansion in areas undergoing the most dramatic folding (Fig. 3), consistent with previous reports in ferret (16, 20). To quantify change in preterm folding, we analyzed non-dimensional mean curvature, defined as *K* = KL* where 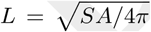 and *SA* = total cortical surface area (20). As shown in Fig. 1*B* (right), *K\** = 1 across a spherical surface, and the global (or local) average of *|K*|* increases for more convoluted surfaces. Local folding was estimated as the difference between *|K*|* on the older and younger surfaces after one iteration of Gaussian kernel smoothing (*σ* = 4 mm). As shown in Fig. 4*B*, regions of highest cortical folding were similar, but not identical, to those of highest cortical expansion.

### Cortical growth changes regionally and dynamically during folding

In individual growth measurements calculated from scans less than 6 weeks apart (n=27 measurements from 15 subjects), we also investigated temporal variations in local growth. As shown in Fig. 5*A*, relative expansion initially appears highest in the primary motor, sensory, and visual cortices, as well as the insula, but lower near term-equivalent (n=4 with four sequential scans). By contrast, relative growth appears to increase with age in parietal, temporal, and frontal lobes. To determine whether these dynamic shifts were statistically significant, we considered the effect of midpoint PMA (halfway between younger and older scan PMAs) as a covariate. PALM with threshold-free cluster enhancement revealed mean spatial patterns (Fig. 5*B* left, mean PMA=33 weeks) similar to those for 30 to 38 weeks (Fig. 4*B*, mean PMA=34 weeks). Importantly, temporal analysis revealed significant decreases in relative expansion in the insula and primary motor, sensory, and visual cortices (i.e. initially fast expansion of these regions slows down relative to other regions), as well as increases in the lateral temporal lobe over time (Fig. 5*B*).

**Fig. 5.**
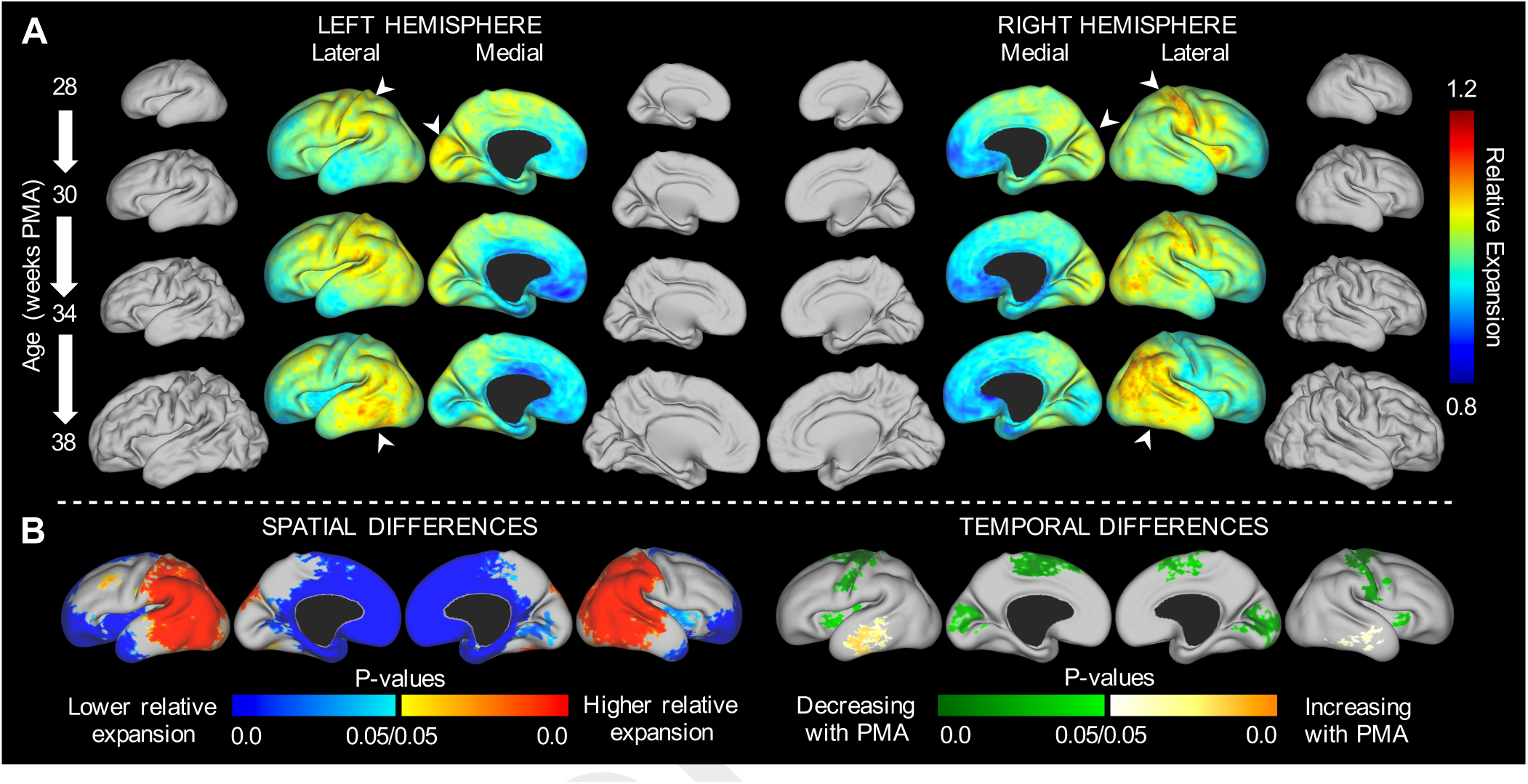
Regions of highest cortical expansion change over time. *(A)* Maps of average relative expansion are shown for short periods of development, denoted on left. For subjects in which three distinct periods of growth could be measured (n=4), regions of maximum expansion (white arrowheads) appear to shift over time. For illustration, average mid-thickness surfaces are also shown to scale for each time point. (B) Statistically significant regional differences were observed based on 27 growth measurements (15 subjects) over the course of the third trimester equivalent (temporal resolution <6 weeks, mean PMA=33 weeks). Left: Relative expansion is higher in the lateral parietal, temporal, occipital and frontal regions (red) and lower in the medial frontal and insular regions (blue). Right: Relative expansion in the primary motor, sensory, and visual cortices, as well as in the insula, decreases over time (green). By contrast, relative expansion increases in the temporal lobe over time (yellow).

We also examined dynamic changes in terms of *growth rate*, defined as local percent increase in cortical area per week. By plotting percentage growth rate versus midpoint PMA at specific locations (Fig. 6*A*), we quantified the rate and acceleration of cortical growth over time. For non-injured individuals (blue and red dots), growth rates were generally higher at earlier ages. Note that growth decelerates significantly (p<0.05) in the initially fast-growing primary cortices (1-5) and insula (8), but it remains fairly constant in the frontal (6-7), temporal (9) and parietal (10) lobes. Linear fit and statistics are available for each region in Table S2.

**Fig. 6.**
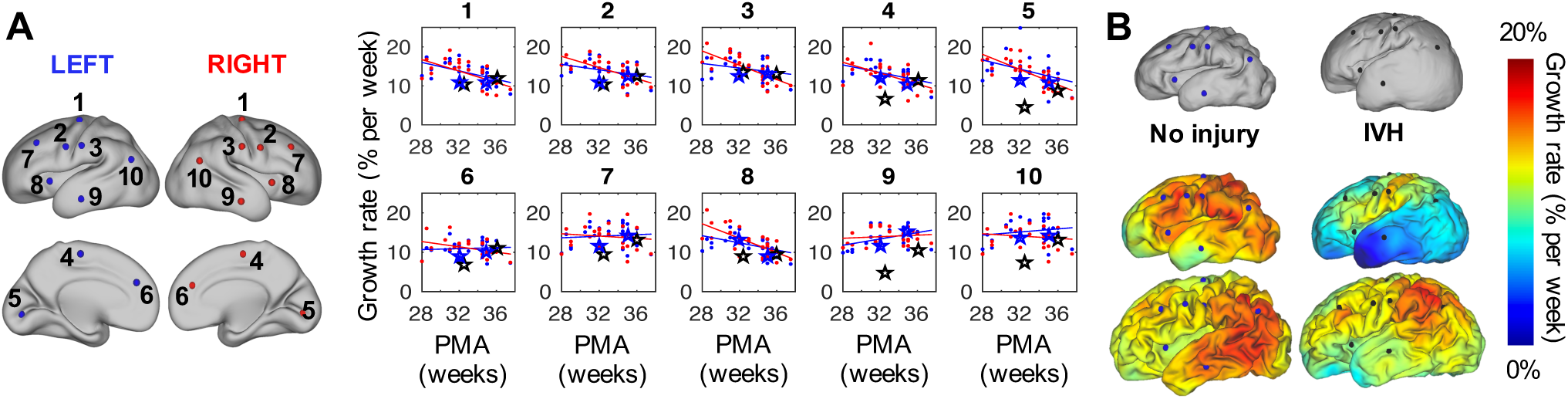
Growth rate decreases in initially fast-growing cortical areas. *(A)* To quantify local changes overtime, local growth rates (n=27 measurements from 15 non-injured subjects) are plotted against midpoint PMA at ten locations on left (blue) and right (red) hemispheres. Growth rate decreases with PMA in locations 1-5 (primary motor, sensory, and visual cortices) and 8 (insula). By contrast, locations 6, 7, 9, and 10 (frontal, temporal, and parietal lobes) remain relatively constant. (*B*) Individual growth rates were also compared between a non-injured subject (left) and a subject with bilateral grade lll/IV intraventricular hemorrhage (IVH, right). From top to bottom, surfaces are shown at approximately 30 weeks, 34 weeks, and 38 weeks PMA. Growth rate from 30 to 34 weeks is plotted on the 34-week surface, and growth rate from 34 to 38 weeks is plotted on 38-week surface. For the subject with IVH, growth rate is initially reduced in occipital (5) and temporal (9) lobes, but later recovers to near non-injured levels. Rates at locations 1-10 in these specific individuals are denoted in (*B*) by blue stars (no injury) or black stars (IVH).

### Local growth estimation detects abnormalities associated with preterm injury

Since our technique produces continuous estimates of cortical growth for individual subjects, we also analyzed cortical surfaces of infants who were identified to have grade III/IV IVH and/or ventriculomegaly during their Neonatal Intensive Care Unit course (n=6). For illustration, Fig. 6*B* compares a subject with no injury (same subject as Fig. 3) to one diagnosed with bilateral grade III/IV IVH. Reduction in growth rate is particularly evident in the temporal and occipital lobes of the subject with IVH (Fig. 6*B*, middle). We also note that growth rate ‘recovered’ to near non-injured levels during the period from 34 to 38 weeks PMA in this individual (Fig. 6*B*, bottom, and black open stars in Fig. 6*A*).

## Discussion

In this work, we implemented an automated, quantitative method for analyzing local growth in longitudinal studies of cortical maturation. Individual registration with aMSM not only produced accurate alignment of gyri and sulci, but it also effectively minimized distortions of the cortical surface (Fig. 3, Table S1). This shift in focus - regularizing the physical anatomical mesh instead of the abstract spherical mesh - offers a significant improvement for longitudinal registration using spherical techniques, which have been plagued by artifactual deformations that obscure real trends and limit interpretation of cortical growth maps (18). Furthermore, our mechanics-inspired regularization penalty (strain energy density, Eq. 1) is physically justified for longitudinal registration and has been shown to outperform other mathematical approaches (18).

Other registration techniques have been proposed to control distortion on the cortical surface via spectral matching or varifolds (21-23), but these have not been integrated into widely-used analysis pipelines (9, 26, 27) or produced smooth, meaningful maps of cortical surface expansion. Furthermore, spherical registration provides an efficient, versatile framework for inter-and intra-subject analysis based on a variety of imaging modalities (17, 28). While this study matched curvatures, an intrinsic feature of any cortical reconstruction, MSM allows registration based on multimodal data, which may further improve the accuracy of registration (9). Future studies may exploit additional data, such as myelin content and fMRI contrasts, to establish correspondence, although such measures are often unavailable in preterm or fetal studies.

The current approach provides continuous maps of cortical expansion for each individual, enabling computation of continuous statistics across the surface (25). Without the limitations of ROI analysis (14-16), spatiotemporal trends presented here offer new insight into the trajectory of cortical growth and maturation. In many respects, these patterns are consistent with past literature. Diffusion tensor imaging and histological analyses have reported mature dendritic branching in the primary motor and sensory cortices before the visual cortex, which in turn matures earlier than the frontal cortex (1, 2). Similar patterns have been reported for regional folding in the preterm brain (7). Our dynamic measures of cortical surface expansion bridge the gap between these metrics, supporting the idea that biological processes (neuronal migration and dendritic branching) produce physical expansion of the cortex (constrained regional growth), which leads to mechanical instability and folding in different areas at different times.

As shown in Fig. 7, the patterns we report for preterm growth (third trimester equivalent) may also link existing studies of cortical growth during the late second trimester and childhood. A volumetric study of fetal MRIs found maximum cortical growth at the central sulcus, which increased from 20-24 weeks to 24-28 weeks, as well as above-average growth near the insula, cingulate, and orbital sulcus (29). These patterns are consistent with our results for 28-30 weeks (Fig. 7*A*, top). Furthermore, we observe a trajectory in which the area of maximum growth migrates outward from the central sulcus, toward the parietal then temporal and frontal lobes, while dissipating from the medial occipital lobe (Fig. 7*C*). As shown in Fig. 7*B*, this is generally consistent with the reported pattern of postnatal expansion (term to adult in human, also proposed across evolution) (10). A limitation of our study is the use of preterm infant data rather than fetal scans, which has become feasible with the advent of improved motion correction tools (30), since studies suggest global surface expansion and folding may develop faster in fetuses (31). While our results appear consistent with second trimester and postnatal trends, future studies should assess potential differences in preterm versus healthy growth as *in utero* longitudinal scans become available with sufficient temporal resolution and sample size.

**Fig. 7.**
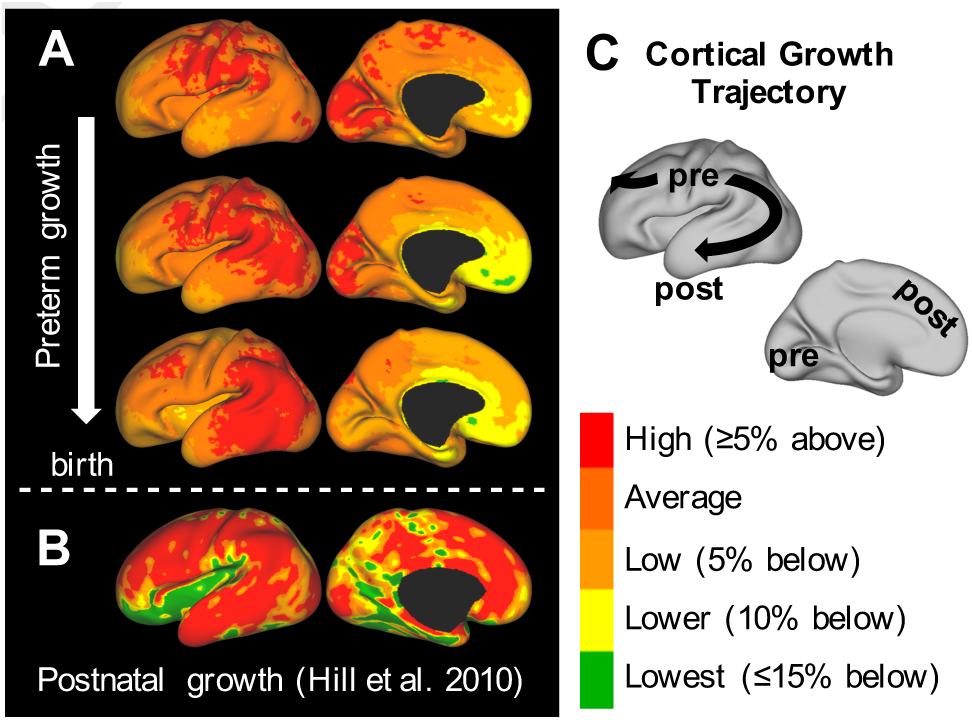
Trajectory of preterm growth may continue after term. *(A)* We observe that local regions of highest expansion (‘hot spots’) migrate smoothly from the central sulcus and nearby regions (top) into parietal (middle) then, finally, frontal and temporal regions (bottom). Growth slows gradually in the primary visual cortex. (*B*) Reported postnatal trends appear to continue this trajectory, maintaining high growth in the parietal, temporal and frontal lobes (10). Expansion is lowest in the insula and visual cortex after birth (though still a minimum of two-fold, adapted from Fig. 1 of ref. (10)). (C) Schematic illustrating the trajectory of the maximum growth region from primary motor, sensory and visual cortices - areas highly conserved across species and critical for basic survival - into regions highly developed in humans, particularly after birth. pre = prenatal/preterm, post = postnatal.

We also note some conflicting results with respect to past studies that analyzed manually-defined ROIs. For example, a recent study investigating the same period of development in preterm infants (30 weeks PMA to term-equivalent) analyzed growth by dividing the brain into its major lobes (14); it reported high expansion in the parietal and occipital lobes, in agreement with our results, but low expansion in the temporal lobe. We speculate that this discrepancy stems from inclusion of the insula and medial temporal surface in their temporal lobe ROI. Relatively low expansion of these regions (Fig. 4*B*) may be sufficient to ‘cancel out’ the high expansion we observed in the lateral temporal region. Another recent study used ROIs to relate regional surface expansion and cellular maturation in the developing ferret brain (16). This ROI analysis did not reveal a clear relationship, whereas our current approach in human (Fig. 6) (1, 2) and ferret (32) suggests a connection between the two. We speculate that *a priori* definition of ROIs may diminish the ability to detect such relationships.

Finally, we note that, for infants with high-grade IVH and/or ventriculomegaly, our analysis was able to clearly detect alterations in cortical growth. Fig. 6*B* provides an example of grade III/IV IVH, where abnormal folding is evident at all time points. However, as illustrated in Fig. 4*B*, folding may not serve as a perfect representation of underlying growth. Our method revealed reduced growth rate in specific regions, followed by recovery to near-normal levels. These exact areas and effects would be difficult to pinpoint with global measures, or even local measures of folding. Just as we were able to detect subtle but consistent differences in our temporal analysis of growth patterns, future studies may be able to detect differences due to specific injury mechanisms, genetic disorders, or environmental variables.

## Materials and Methods

### Recruitment, MRI acquisition and surface generation

Preterm infants used in this study were born at <30 weeks gestation and were recruited from St Louis Children’s Hospital. The Washington University Institutional Review Board approved all procedures related to the study, and parents or legal guardians provided informed, written consent. Images were obtained using a turbo spin echo T2-weighted sequence (repetition time = 8,500 milliseconds; echo time = 160 milliseconds; voxel size = 1 x 1 x 1 mm^3^) on a Siemens (Erlangen, Germany) 3T Trio scanner. T2-weighted images were processed, and cortical segmentations were generated at the mid-thickness of the cortex using previously documented methods (33). These segmentations were then used to generate cortical surface reconstructions, including mid-thickness, inflated, flat, and spherical surfaces, for each hemisphere using methods previously reported (33). Preterm infants with moderate to severe cerebellar hemorrhage, grade III/IV IVH, cystic periventricular leukomalacia or ventriculomegaly on MRI were identified and analyzed separately (34, 35). A summary of clinical and demographic information for included and excluded subjects can be found in Table S3.

### Longitudinal surface alignment and theory

To obtain point correspondence between individual surfaces across time, cortical surfaces (‘input’ and ‘reference’) were projected to a sphere and registered with aMSM (18) (Fig. 2). This tool systematically moves points on the input spherical surface in order to maximize similarity of a specified metric and minimize strain energy between the *anatomical* input and reference surfaces. Mean curvatures, generated in Connectome Workbench (24), were used for matching data and cortical mid-thickness surfaces, rescaled to the same total surface area, were used as the anatomical input and reference surfaces.

Inspired by studies that have modeled brain tissue as a hyperelastic material (3, 4, 6, 36), we define surface strain energy as:

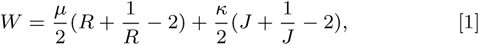

where *R = λ*_1_*/λ*_2_ represents change in shape, *J* = *λ*_1_*λ*_2_ represents change in size, and *λ*_1_ and *λ*_2_ represent in-plane anatomical stretches in the maximum and minimum principal directions, respectively. This form corresponds to a modified, compressible Neo-hookean material in 2D, and here we define bulk modulus, re, to be 10 times greater than shear modulus, *μ.* To prevent bias associated with the direction of registration (37), average results were calculated for aMSM registration performed from both older to younger and younger to older surfaces (Fig. S3). Additional details on theory and implementation of aMSM, as well as parameter effects of bulk-to-shear ratio, are described in *SI text* and (18). Other parameter effects have been described previously (17, 18).

### Group statistics

To analyze trends in growth, individual metric maps were compared on the 30-week group atlas by applying point correspondences described in Fig. 4. Atlas generation details can be found in *SI text.* Individual-to-atlas alignment was accomplished with aMSM, using one surface from each individual (time point closest to 30 weeks PMA) as input. Permutation Analysis of Linear Methods (PALM) was performed with threshold-free cluster enhancement (TFCE) (25), using 1000 iterations and a medial wall mask. Single group t-tests were performed on the log transform of relative surface expansion to obtain a normally distributed metric centered at zero. Mean 30-week mid-thickness surfaces and vertex areas were used for TFCE surface area computations.

Significance of correlations was assessed using Pearson’s correlation coefficient in MATLAB, and total strain energy was calculated by integrating Eq. 1 with respect to cortical surface area (The MathWorks, Inc., Natick, MA) (19). Significant improvements due to aMSM were assessed by comparing total correlation and energy values before and after registration with paired t-tests (Table S1). For local analysis of growth rate (Fig. 6), one iteration of Gaussian kernel smoothing *(σ =* 2 mm) was applied (24) before determining linear fit and correlation values in MATLAB. Similar linear fits were obtained with no smoothing and with increased smoothing.

## ACKNOWLEDGMENTS

This work was supported by NIH grants R01 NS055951 (PVB), R01 NS070918 (LAT), T32 EB018266 (KEG), K02 NS089852 (CDS), K23 MH105179 (CER), UL1 TR000448 (CER and CDS), 5U54HD087011 (DLD), and R01 HD057098 (cohort); and the Intellectual and Developmental Disabilities Research Center at Washington University (P30 HD062171). We gratefully acknowledge Jeff Neil, Joe Ackerman, Jr., Karen Lukas and Anthony Barton for their contributions to this study.

